# Bispecific antibodies against the hepatitis C virus E1E2 envelope glycoprotein

**DOI:** 10.1101/2024.10.04.616615

**Authors:** Laura Radić, Anna Offersgaard, Tereza Kadavá, Ian Zon, Joan Capella-Pujol, Fabian Mulder, Sylvie Koekkoek, Vera Spek, Ana Chumbe, Jens Bukh, Marit J van Gils, Rogier W Sanders, Victor C Yin, Albert J R Heck, Judith M Gottwein, Kwinten Sliepen, Janke Schinkel

**Affiliations:** Department of Medical Microbiology and Infection Prevention, Amsterdam Infection & Immunity Institute, Amsterdam UMC, University of Amsterdam, Meibergdreef 9, 1105AZ Amsterdam, The Netherlands; Amsterdam Institute for Immunology and Infectious diseases, Amsterdam, the Netherlands; Department of Microbiology and Immunology, Weill Medical College of Cornell University, 1300 York Ave, New York, NY 10065, United States; Copenhagen Hepatitis C Program (CO-HEP), Department of Infectious Diseases, Copenhagen University Hospital–Hvidovre, 2650 Hvidovre, Denmark, and Department of Immunology and Microbiology, Faculty of Health and Medical Sciences, University of Copenhagen, 2200 Copenhagen N, Denmark; Biomolecular Mass Spectrometry and Proteomics, Bijvoet Center for Biomolecular Research and Utrecht Institute for Pharmaceutical Sciences, Utrecht University, 3584 CH Utrecht, the Netherlands; Netherlands Proteomics Center, 3584 CH Utrecht, the Netherlands

**Keywords:** hepatitis C virus (HCV), bNAbs, bispecific antibodies, E1E2 envelope glycoprotein, neutralization, breadth, avidity, mass photometry

## Abstract

Hepatitis C virus (HCV) currently causes about one million infections and 240,000 deaths worldwide each year. To reach the goal set by the World Health Organization (WHO) of global HCV elimination by 2030, it is critical to develop a prophylactic vaccine. Broadly neutralizing antibodies (bNAbs) target the E1E2 envelope glycoproteins on the viral surface, can neutralize a broad range of the highly diverse circulating HCV strains and are essential tools to inform vaccine design. However, bNAbs targeting a single E1E2 epitope might be limited in neutralization breadth, which can be enhanced by using combinations of bNAbs that target different envelope epitopes. We have generated 60 IgG-like bispecific antibodies (bsAbs) that can simultaneously target two distinct epitopes on E1E2. We combine non-overlapping E1E2 specificities into three types of bsAbs, each containing a different hinge length. The bsAbs show retained or increased potency and breadth against a diverse panel of HCV pseudoparticles (HCVpp) and HCV produced in cell culture (HCVcc) compared to monospecific and cocktail controls. Additionally, we demonstrate that changes in the hinge length of bsAbs can alter the binding stoichiometry to E1E2. These results provide insights into the binding modes and the role of avidity in bivalent targeting of diverse E1E2 epitopes, and suggest structural differences between HCVpp and HCVcc. This study illustrates how potential cooperative effects of HCV bNAbs can be utilized by strategically designing bispecific constructs. These new HCV bsAbs can guide vaccine development and unlock novel therapeutic and prophylactic strategies against HCV and other (flavi)viruses.

## Introduction

Hepatitis C virus (HCV), a member of the *Flaviviridae* family, is a bloodborne virus that frequently leads to chronic hepatitis, which can progress to liver cirrhosis and cancer after years of indolent infection. Globally, an estimated 50 million people live with hepatitis C, with around one million new infections and approximately 240000 deaths annually (1). Although direct-acting antivirals (DAAs) can cure HCV in >95% of cases, access to treatment remains limited for many and those cured are susceptible to reinfections (2, 3). In some cases, multi-DAA resistance can occur, and there are often no treatment options for those who failed multiple treatment regimens (4, 5). Therefore, the development of effective vaccines is essential to curb the ongoing epidemic (6, 7).

One of the major obstacles in eliminating HCV is its high sequence diversity - approximately 30% between the major HCV genotypes, with genotypes 1-3 accounting for 80% of infections (8). This diversity exceeds that of most viruses, including the notably diverse human immunodeficiency virus 1 (HIV-1) (7, 9, 10). Fortunately, a proportion of HCV-infected individuals spontaneously recover, which has been linked to the development of broadly neutralizing antibodies (bNAbs) capable of neutralizing a wide range of circulating variants (11–16). Passive administration of HCV bNAbs in animal models has been shown to protect against (re)infection, but also help clear ongoing HCV infection (17–19). Potent bNAbs hold therapeutic potential in humans as long-lasting protection in the absence of a vaccine or as an intervention method (20).

HCV bNAbs target overlapping but distinct epitopes on the E1E2 envelope glycoprotein complex (16, 21–25). E1E2, anchored into the viral membrane, consists of E2, which interacts with several cellular receptors such as CD81 and scavenger receptor class B type I (SR-B1), and E1, of which the function is less clear, but it is thought to be involved in membrane fusion and virus assembly (26, 27). Although some E1E2-targeting bNAbs can neutralize multiple viral strains, it is unlikely that a single bNAb will be able to potently neutralize all strains. To prevent viral escape, a promising approach is to use combinations of bNAbs. By selecting bNAbs that cooperate or act synergistically when binding to their distinct epitopes, it is possible to not only improve neutralization breadth but also increase overall potency, for example by using a bNAb cocktail (18, 28, 29).

Another strategy to enhance antibody neutralization is by combining different bNAb specificities into multivalent reagents, such as bispecific antibodies (bsAbs) (30–33). While conventional Abs have two fragment antigen-binding (Fab) arms that bind the same epitope of an antigen, bsAbs are engineered to have two different Fabs, each capable of simultaneously targeting either different epitopes on the same antigen or two different antigens. Several antiviral bsAbs have proved to be successful in *in vitro* and *in vivo* preclinical studies, in particular against SARS-CoV-2, and bi- and tri-specific HIV-1 bNAbs are currently being tested in clinical trials (31–35). However, to our knowledge, bsAbs against HCV E1E2 have not yet been described.

Here, we produced and characterized a panel of 60 bsAbs, each in one of three different formats with varying hinge length, that simultaneously target two distinct E1E2 epitopes. In neutralization assays against HCV pseudoparticles (HCVpp) and HCV produced in cell culture (HCVcc) of strains from different genotypes, several bsAbs outperformed their parental counterparts and corresponding cocktails, with respect to the neutralization breadth and potency.

Using single molecule mass photometry, we compared the binding stoichiometry of the bsAbs to their parental bNAbs revealing how hinge length influences their mode of action. By combining antibody specificities into bispecific constructs, it is possible to expand the range of binding modes beyond what is achievable with conventional HCV bNAbs. This proof-of-concept study demonstrates how existing HCV bNAbs can be improved through cooperativity in novel bsAb formulations.

## Results

### Design and characterization of HCV E1E2-targeting bispecific antibodies

For the generation of bispecific antibodies, we chose bNAbs targeting different epitopes of E1E2. Informed by the recently described structure of the HCV E1E2 envelope glycoprotein (36, 37) and previous data on antibody specificities we defined different combinations of bNAbs with a promising potential for synergistic or complementary activity when used in a bispecific format. These combinations included AR4A and AT1618, which target a conformational epitope on E2 (antigenic region 4 (AR4)) and require the full native(- like) E1E2 complex for binding (36, 38). Further, AR3C, HEPC74 and AT1209, which bind to epitopes overlapping with the binding site of CD81 (CD81bs) on E2, and HC84.26, which binds to antigenic site 434, at the front layer of the CD81bs (Fig. 1A). Finally, AP33 which binds a linear epitope on E2 and IGH505 which targets E1.

**Figure 1.**
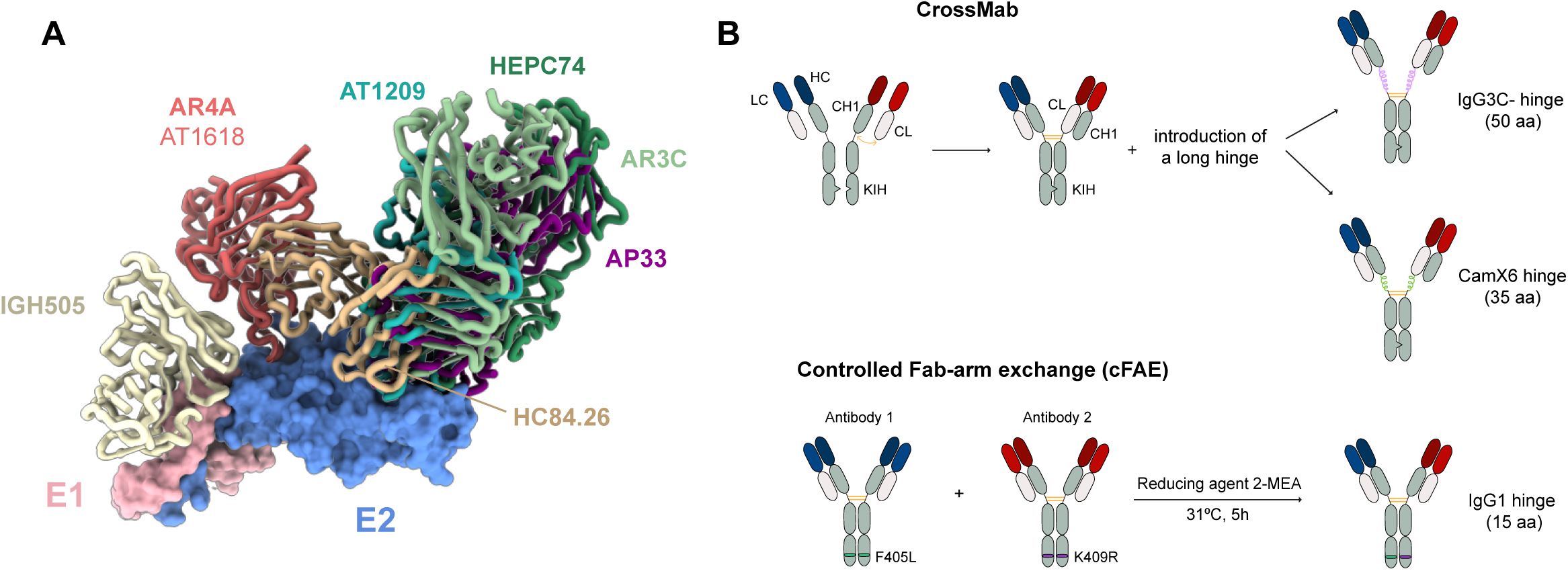
Design of three types of HCV E1E2-targeting bispecific antibodies. **A** Structural representation of the E1E2 glycoprotein in complex with HCV bNAbs AR4A, IGH505, AT1209 (PDB: 7T6X), AR3C (PDB: 4MWF), HEPC74 (PDB: 6MEH), AP33 (PDB: 4G6A) and HC84.26 (PDB: 8THZ). The structure of AT1618 is not available, The structure of AT1618 is not available, but in alanine scanning mutagenesis epitope partly overlaps with that of AR4A and its binding depends on E1E2 (16). **B** Schematic depiction of the methods of bsAb generation used in this study: CrossMab formats with an IgG3C- or Camelid (CamX6) hinge and controlled Fab-arm exchange. HC, heavy chain; LC, light chain; CH1, constant heavy chain region 1; CL, constant light chain region; KIH, knob-in-hole; aa, amino acid.

Previous studies demonstrated that the length of the hinge between the fragment crystallizable (Fc) and Fab domains can greatly affect the breadth and potency of bsAbs (30, 39). Therefore, we combined pairs of bNAbs targeting distinct epitopes and generated three types of bsAbs with varying hinge lengths, as detailed in the method section (Fig. 1B). The bsAbs with longer hinges were produced in a CrossMab IgG1 format, with either a 50 amino acid hinge, derived from an IgG3 hinge (with the first 9 of 11 cysteines removed to enhance Fab flexibility, referred to as IgG3C-hinge) (30) or a 35 amino acid hinge, from camelid heavy chain antibody constructs (CamX6 hinge) (40). Supplementary fig. 1 details the modifications in elbow sequences between the CH1 and CH2/CL regions and the hinge sequences. To generate bsAbs with a 15 amino acid “normal” IgG1 hinge (referred to as “IgG1 hinge bsAbs”), we used controlled Fab-arm exchange (cFAE) (41) (Fig. 1B). In this method matching point mutations (F405L/K409R) are introduced in the Fc regions of the parental antibodies, enabling Fab-arm exchange under reducing conditions resulting in bsAb formation. In total, we designed 21 different combinations in the three bsAb formats with different hinge lengths for a total of 63 bsAb designs. We managed to successfully produce 60 of these.

The bsAbs showed distinct bands for both the heavy and light chains on reducing SDS-PAGE, with a noticeable increase in heavy chain size for formats with greater hinge lengths (Fig. 2A, Suppl. fig. 2). To confirm whether the bsAbs contained the two separate arms as designed, we performed an enzyme-linked immunosorbent assay (ELISA). We tested binding of recombinant E2 to the AR4A/HC84.26 bsAbs, using the individual AR4A and HC84.26 monoclonal antibodies (mAbs) and the corresponding antibody cocktail as controls. HC84.26 binds recombinant E2, while AR4A does not, since its binding requires the full native(- like) E1E2. Antibody binding was detected using secondary antibodies against κ or λ light chain isotypes, which are specific to AR4A and HC84.26, respectively (Fig. 2B). As expected for the individual mAbs, we detected a signal only for HC84.26 (i.e. λ chain), but not for AR4A (i.e. κ chain). Similarly, for the cocktail of AR4A and HC84.26, we only observed a signal when using the anti-λ secondary antibody, stemming from the binding of HC84.26. In contrast, all three AR4A/HC84.26 bsAbs showed signals for both the κ and λ chains, confirming that both arms were present in a single bsAb molecule. Representative bsAbs maintained their thermostability, as determined by differential scanning fluorimetry (DSF) with melting temperatures in the 70-75°C range, comparable to monospecific bNAb controls (Fig. 2C, D). We also wanted to confirm that the mutations introduced into the Fc regions to generate our bsAbs did not affect Fc effector functions, such as antibody-dependent cellular cytotoxicity (ADCC). Natural killer cells recognize the Fc region of antibodies via Fc gamma receptor IIIa (FcγRIIIa), triggering apoptosis through the release of cytotoxic granules (42). In our ELISA experiments, representative AR4A/AR3C bsAbs of all three formats showed no impairment in binding to soluble E1E2, nor in subsequent engagement by FcγRIIIa (Fig. 2E).

**Figure 2.**
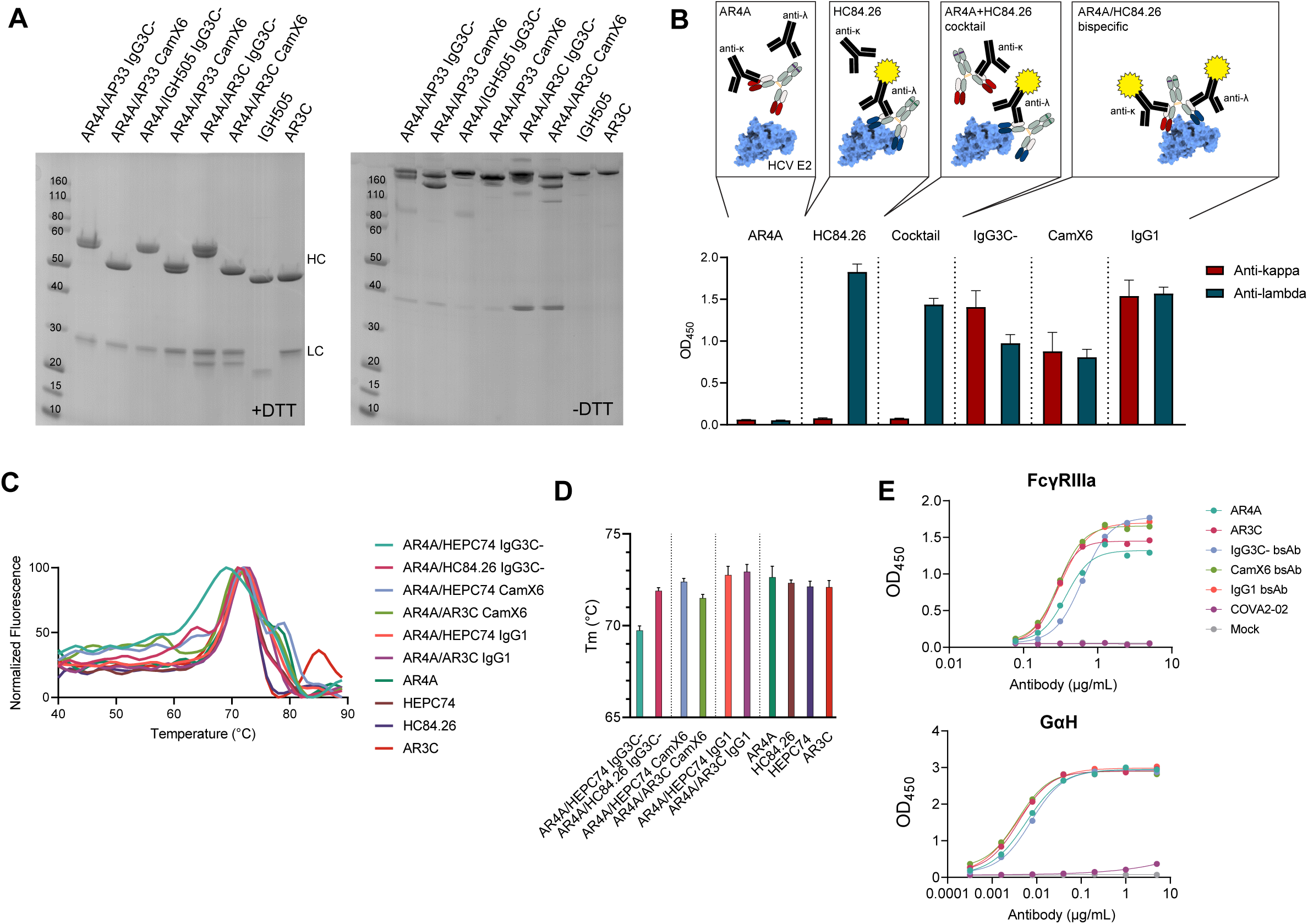
Bispecific antibodies are well folded, thermostable and contain two distinct Fab arms. **A** SDS-PAGE gels under reducing (left) and non-reducing (right) conditions of representative CrossMab bsAbs with an IgG3C- or CamX6 hinge. MAbs IGH505 and AR3C are used as controls. Heavy chains (HC) and light chains (LC) are labeled at detected sizes. **B** ELISA using LC-isotype specific secondary antibodies. His-tagged HCV E2 was bound to a nickel–nitrilotriacetic acid (NiNTA) ELISA plate, followed by AR4A or HC84.26 mAbs, 1:1 cocktail of AR4A and HC84.26 or an AR4A/HC84.26 bsAb with IgG1, IgG3C- or CamX6 hinge. Signals detected by secondary Abs specific for either kappa (i.e. AR4A) or lambda LC (i.e. HC84.26) are depicted in yellow. **C** Normalized thermal unfolding curves (dF/dT) of selected bsAbs and parental mAbs obtained by differential Scanning Fluorimetry (DSF). Lines represent the mean of triplicate fluorescence intensity measurements. **D** Bar graph of melting temperatures (Tm) of selected bsAbs and parental mAbs. **E** ELISA to test binding of representative AR4A/AR3C bsAbs of each type to FcγRIIIa. Soluble HCV E1E2 was coated on a Strep-TactinXT plate, followed by incubation with the bsAbs, mAb or cocktail controls and soluble FcγRIIIa-dimer. Signal was detected using a streptavidin-HRP detection Ab. A control ELISA (bottom panel) was also performed to detect binding directly after E1E2/antibody incubation using a GαH secondary Ab.

### HCV bsAbs can improve neutralization potency and breadth

To evaluate the neutralization activity of the bsAbs, we performed neutralization assays using HCVpps from three different strains: AMS0232 (genotype 1), UKNP2.4.1 (genotype 2) and UKNP3.2.2 (genotype 3). These strains were selected from a panel of HCVpps (43) for their mid-tier resistance to neutralization and distinct genotypes, allowing us to observe potential differences in neutralization by the different bsAbs. We screened all 60 bsAbs and corresponding cocktails using 2 concentrations of total antibody (0.2 and 2 μg/mL) (Fig. 3A). When comparing the overall antibody categories, no significant differences in neutralization potency were observed (Fig. 3B), although combinations (bsAbs and cocktails) showed a slight trend towards improved potency. Neutralization profiles of the three bsAb types and antibody cocktails correlated well, indicating that for most bsAbs combining two specificities did not lead to enhanced neutralization through synergy between the arms (Suppl. fig 3). However, there were notable exceptions where specific bsAbs demonstrated increased potency compared to their parental mAbs and cocktails (Fig. 3A), such as for IgG1 hinge AR4A/AT1209 bsAb. Interestingly, the hinge length appeared to influence bsAbs potency, e.g. AR4A/AT1209 was less potent with an IgG3-hinge. These results suggest that binding cooperativity of the two Fab arms facilitates synergistic neutralization. Based on these findings, seven of the most interesting candidates were selected for further evaluation and more detailed analysis of their neutralization activity (Fig. 3C).

**Figure 3.**
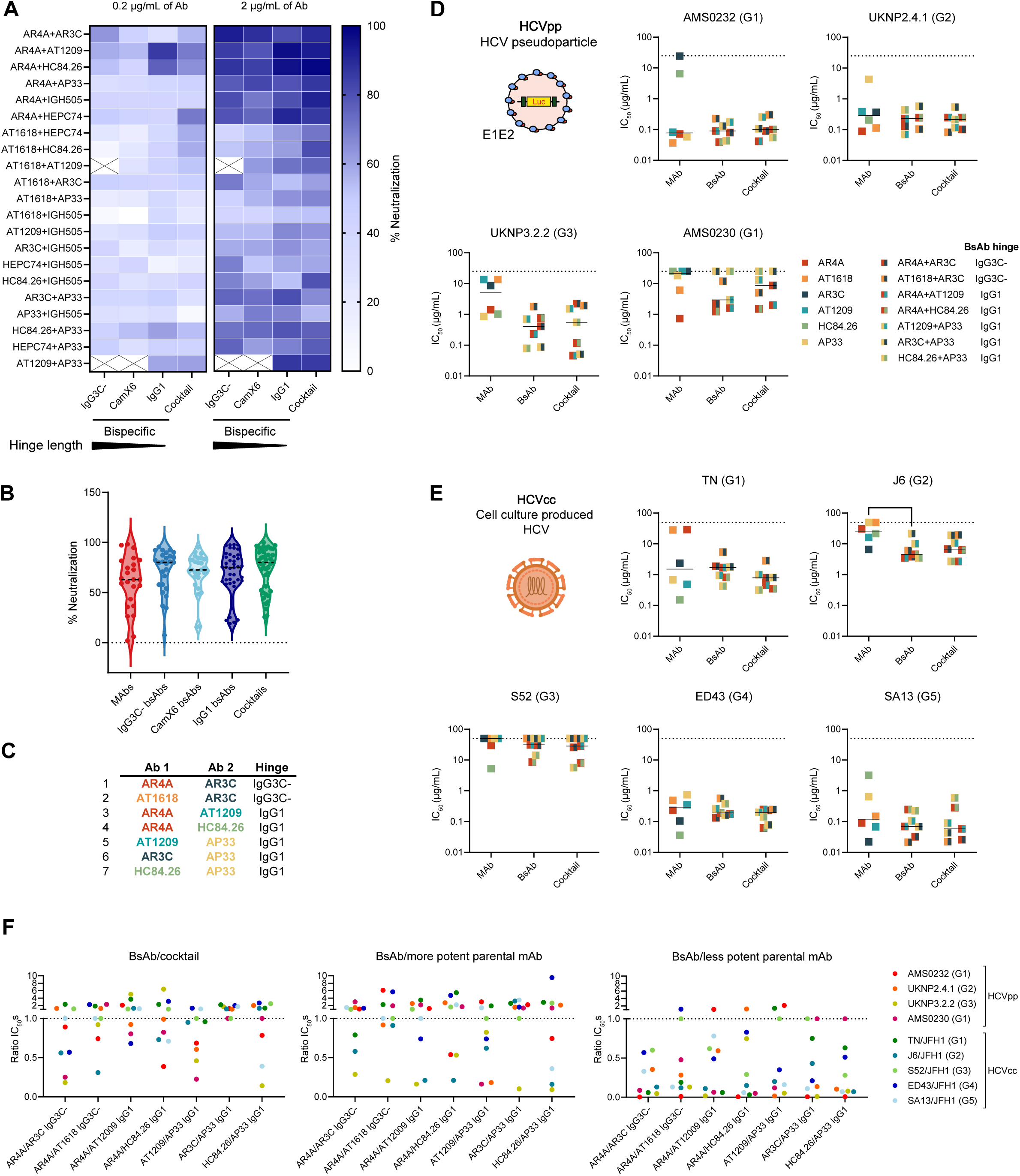
HCV bsAbs display neutralizing activity in HCVpp and HCVcc assay systems. **A** Screen of HCVpp neutralization by 0.2 or 2 µg/mL of total Ab. Heatmap shows percentages of neutralization by the Abs (geomean) obtained from neutralization of 3 different HCVpps (AMS0232, UKNP2.4.1 and UKNP3.2.2). Each heatmap row represents a specific combination of antibody specificities, while each column represents the type of antibody (either of the three bsAb types or a cocktail of mAbs). **B** Percent neutralization of HCVpps by all individual mAbs, bsAbs at 2 µg/mL or cocktails of 2 mAbs at 2 µg/mL total concentration (1 µg/mL of each mAb). Each symbol indicates neutralization of an HCVpp, measured in duplicate. Dotted lines indicate the median. **C** List of seven bsAb candidates selected for further analysis. They were tested against 4 HCVpps (**D**) and 5 HCVccs (**E**). Viral strains of the envelope glycoproteins are listed in the graph titles, with the respective viral genotype in brackets. Each square symbol represents a half-maximal neutralization (IC_50_) value (µg/mL) of neutralization by a mAb (square of one color), bsAb or cocktail (square of 2 colors representing the combination of mAbs), measured in triplicate. Abs with IC_50_s above 25 µg/mL (HCVpp) or 50 µg/mL (HCVcc) were considered non-neutralizing (dotted line). Kruskal-Wallis test, *=p<0.05. When not indicated above the graph, the statistical differences between groups were not significant. **F** IC_50_ ratios between tested bsAb and corresponding cocktail, bsAb and more potent parental mAb and bsAb and less potent parental mAb, for all seven tested antibody combinations, against every tested HCVpp and HCVcc. Ratio <1 represents improved activity of the bsAb against a particular viral strain.

Among the candidates containing the IgG3C-hinge, AR4A/AR3C stood out for its highest potency. To assess the influence of changing specificity without changing the target epitope of one of the arms, we also selected AT1618/AR3C IgG3C-. We included combinations of AR4A with AT1209 and HC84.26, because these showed the highest potency among the IgG1 hinge bsAbs. Additionally, there was potential for improved activity for IgG1 hinge bsAbs pairing AT1209, AR3C or HC84.26 (all targeting epitopes overlapping with or near the CD81bs) with AP33. We next performed neutralization assays with the seven best bsAb candidates, mAbs and cocktail controls, using full concentration-response curves against four HCVpps from different genotypes (Fig. 3D, Suppl. fig. 4A). In addition to the three genotype 1-3 HCVpps used in the initial screen, we included the highly neutralization-resistant genotype 1 strain AMS0230 (43). Overall, combinations of bsAbs and cocktails exhibited greater neutralization activity compared to monospecific antibodies. Notably, all seven bsAbs showed neutralizing activity against every HCVpp, except for two (AR3C/AP33 IgG1 and HC84.26/AP33 IgG1), that did not neutralize AMS0230, which is consistent with the behavior of the parental mAbs. In some instances, the bsAbs showed enhanced neutralization, presumably due to connectivity of the two distinct arms into one molecule and cooperative binding to E1E2. For example, AR4A/HC84.26 IgG1, AT1209/AP33 IgG1 and HC84.26/AP33 IgG1 showed higher potency than both individual mAbs and their corresponding cocktails against AMS0232, UKNP2.4.1 and UKNP3.2.2, respectively (Fig. 3D, Suppl. fig. 4A).

To further characterize the neutralization activity of the seven bsAb candidates against replication-competent virus, we tested them against HCVcc. HCVcc more closely resembles authentic HCV than HCVpp, as it incorporates human apolipoproteins on its envelope and undergoes the full virus life cycle (44). We tested the bsAbs along with mAbs and cocktail controls, against five different HCV recombinants with the structural proteins of strains TN (genotype 1), J6 (genotype 2), S52 (genotype 3), ED43 (genotype 4) and SA13 (genotype 5). All seven bsAbs could neutralize the majority of tested viruses, which ranged from highly sensitive (SA13) to highly resistant (J6 and S52) to antibody neutralization (45) (Fig. 3E, Suppl. fig. 4A). Neutralization was not observed against the resistant strain S52 when both parental mAbs were non-neutralizing, similar to the lack of neutralization seen with HCVpp AMS0230.

Similar to HCVpp, no significant differences in neutralization potency were observed between antibody groups for most HCVcc strains, except for J6, where the bispecific antibodies outperformed the monospecific (Fig. 3E). As expected, most bsAbs exhibited neutralization potencies that fell between those of their parental mAbs. However, in some cases, bsAbs had lower IC_50_ values compared to both parental mAbs and corresponding cocktails, pointing at synergistic neutralization activity. This was observed for AR4A/AR3C IgG3C- and AR4A/AT1209 IgG1 against J6 and HC84.26/AP33 against SA13. In addition to enhancing neutralization potency, bsAbs can also broaden neutralization activity. Overall, bsAbs containing an HC84.26 arm, i.e. AR4A/HC84.26 IgG1 and HC84.26/AP33 IgG1, displayed substantial broad neutralizing activity, even when mAb of the other arm did not show 50% neutralization at the highest tested concentration, as seen with HC84.26/AP33 against S52. This increase in breadth is an important advantage in combating viral escape.

To evaluate potential synergistic activity of the bsAbs, we calculated IC_50_ ratios between the bsAbs and cocktails, and each parental mAb individually, for all tested HCVpp and HCVcc strains (Fig. 3F). All bsAbs had a ratio lower than one against at least one tested virus indicating enhanced potency compared to either the cocktail or the more potent of the parental mAb. The only exception was IgG1 hinge AR3C/AP33 bsAb which showed equal or decreased activity compared to the cocktail. These results suggest that cooperativity between two HCV mAbs can be achieved when combined into a bispecific format, while also enhancing neutralization breadth. When comparing the neutralization activity of the bsAbs with the less potent parental mAb, we observed a reduced potency for the bsAb only in a few cases, primarily when both parental mAbs were non-neutralizing against the respective viral strain. In summary, neutralization activity is not compromised by creating bsAbs. In fact, bsAbs often surpass the neutralization potency of the less potent parental mAb and in many cases also the more potent parental mAb. The latter observation indicates that bsAbs can achieve synergy by binding to both epitopes simultaneously.

To identify the bsAb with the highest overall breadth, potency and enhancement over the parental mAbs and cocktail, we performed a statistical ranking analysis, comparing all tested antibodies on their IC_50_, the ratio of bsAb and cocktail, and ratios of bsAb to each parental mAb individually (Suppl. fig. 4B). Interestingly, the ranking suggested that bsAbs with the long IgG3-hinges were more effective at neutralizing HCVcc, while those with IgG1 hinges were superior against HCVpp. In particular, AR4A/AR3C IgG3C-emerged as the best performing candidate against HCVcc.

### Increased bsAb hinge length influences binding stoichiometry

To examine the role of the hinge length in function and binding, we studied the AR4A/AR3C combination in the three bsAbs formats in greater detail. When bound to their respective epitopes, AR4A and AR3C Fabs are 75Å apart at the C-terminal ends of their heavy chains (Fig. 4A). The usual span distance of an IgG1 is 100-150Å (46, 47). We thus hypothesized that 75Å might be too close for simultaneous binding by a bsAb with a short and rigid IgG1 hinge. However, a bsAb with a long and flexible open IgG3C-hinge may be able to bridge these epitopes and allow dual binding on a single E1E2 complex (30).

**Figure 4.**
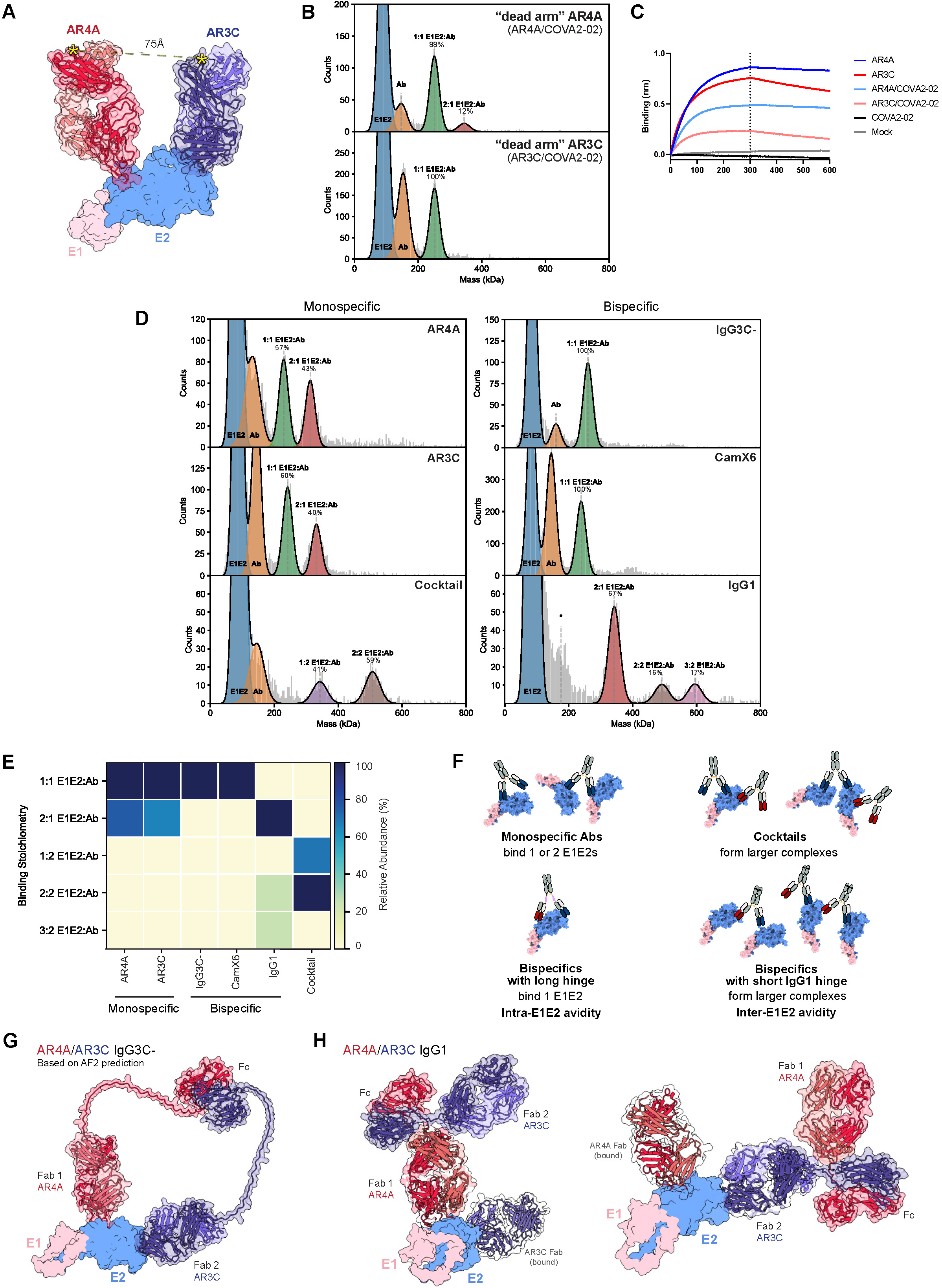
Hinge lenght influences the avidity and stoichiometry of AR4A/AR3C bsAbs binding to E1E2. **A** Fabs of AR4A (PDB: 8FSJ) and AR3C (PDB: 4MWF) in complex with E1E2 (PDB:7T6X), showing the distance between the C-terminal ends of their respective heavy chains (indicated with yellow asterisks). **B** Mass photometry measurements of “dead arm” AR4A and AR3C bsAbs in complex with soluble E1E2 protein. The resulting mass histograms were fitted with Gaussian curves to represent the distinct assemblies. All measurements were performed with a 3:1 E1E2:Ab ratio. **C** BLI traces showing the binding of AR4A, AR3C and “dead arm” bsAbs to E1E2. The SARS-CoV-2 mAb COVA2-02 was used as a binding control, confirming that the “dead arm” bsAbs had one arm that was inactive for E1E2 binding. **D** Mass photometry measurements of mAbs AR4A, AR3C, bsAbs with varying hinge lengths, and AR4A/AR3C cocktail in complex with soluble E1E2 protein. An asterisk indicates overlapping distribution of the IgG1 bsAb with E1E2 contamination (Suppl. fig. 5). **E** Binding stoichiometries observed by MP are summarized as relative abundance (%) of measured E1E2:Ab complexes for the different mAbs, bsAbs and the cocktail. **F** Schematic representation of preferred binding stoichiometries and avidity effects for a representative mAb, bsAbs with short and long hinges and a cocktail of mAbs. **G** Schematic view based on AlphaFold2 prediction of AR4A/AR3C IgG3C-binding, superimposed on the available E1E2 structure (PDB: 7T6X). The AR4A arm of the bsAb is colored in red, the AR3C arm in blue. Darker shades represent the heavy chains, while lighted shades represent the light chains. **H** One Fab arm of a human IgG1 antibody (PDB: 1IGY) aligned with the predicted AR4A Fab (left panel) or AR3C Fab (right panel), colored the same way as in A, to indicate that the angle between the twoFabs of the AR4A/AR3C IgG1 hinge bsAb does not allow simultaneous binding of both epitopes. The other Fab (AR3C in the left panel, AR4A in the right) is shown bound to its epitope, and with a transparent surface.

We utilized mass photometry (MP), a technique capable of precisely measuring the molecular weight distributions of highly heterogeneous biomolecular mixtures through single-molecule light scattering (48). This allows for accurate identification of different binding stoichiometries by their differences in mass (49). We previously used MP to determine the preferential binding modes of mAbs and bsAbs to the trimeric SARS-CoV-2 spike protein (32), distinguishing between different types of binding avidity. We here aimed to determine binding stoichiometries of the HCV bsAbs with different hinges to soluble E1E2, to assess their preferred binding modes. First, we measured the mass distributions for the antibodies and E1E2 separately. As expected, we observed peaks corresponding to ∼150 kDa for all mAbs and IgG1 hinge bsAbs. The longer hinges resulted in increased masses: ∼160 kDa for AR4A/AR3C CamX6 and ∼168 kDa for AR4A/AR3C IgG3C-. For E1E2, a mass of ∼88 kDa was determined (Suppl. fig. 5).

To validate that we can accurately recognize E1E2:antibody (E1E2:Ab) complexes by MP after incubation, we measured complex formation of E1E2 and “dead arm” bsAbs, generated by cFAE. These bsAbs have one functional arm, either AR4A or AR3C, while the other arm is an irrelevant arm of SARS-CoV-2 specific mAb COVA2-02 (50). Functionally, these constructs behave like monovalent Fabs of AR4A and AR3C. The MP histograms revealed populations of unbound E1E2, unbound antibody and E1E2:Ab complexes (Fig. 4B). The mass histograms of the E1E2:Ab complexes translate to particular binding preferences of the antibody to E1E2. As expected, both “dead arm” bsAbs bound E1E2 with a 1:1 stoichiometry (Fig. 4B). The low fraction (∼12%) of 2:1 E1E2:Ab binding observed for “dead arm” AR4A possibly stems from a small proportion of AR4A molecules that did not undergo the cFAE process and remained bivalent. This is consistent with the typical cFAE process efficiency of ∼90-95% (51).

To assess whether removing one functional Fab arm affects binding affinity, we performed a BLI run with both mAbs and “dead arm” bsAbs. As expected, we observed a loss of binding for AR4A and AR3C when paired with an irrelevant arm, indicating that these mAbs require both arms to be functional to strongly bind to their epitopes (Fig. 4C). This finding emphasizes the contribution of avidity in the bsAb constructs, as combining AR4A and AR3C showed restored binding strenght compared to the single mAbs in ELISA (Fig. 2E). Moreover, in most cases we observe retained or even increased neutralizing activity of the bsAbs (Fig. 3A, D, E).

Next, we incubated monospecific bivalent AR4A and AR3C antibodies with three-fold molar excess of E1E2. Both AR3C and AR4A exhibited binding in two distinct stoichiometries: 1:1 (∼60%) and 2:1 (∼40%) E1E2:Ab ratios, demonstrating that each monospecific mAb can use either one or both arms to bind one or two E1E2s (Fig. 4D). In contrast, a cocktail of AR3C and AR4A showed peaks corresponding to 1:2 and 2:2 E1E2:Ab complexes, indicating that a mixture of these two monospecific mAbs, which target non-overlapping epitopes, can bind monovalently to one or two E1E2s or drive cross-linking of two E1E2 heterodimers. Similarly, when measuring AR4A/AR3C IgG1 bsAb, we observed higher-order complexes of 2:2 and 3:2 E1E2:Ab ratios, but with a significant proportion (∼67%) of 2:1 bound E1E2s. This indicates that this bsAb favors inter-E1E2 avidity, i.e. cross-linking different E1E2 molecules. While individual mAbs are also able to adopt this mode of binding, they predominantly bind to a single E1E2. Interestingly, when measuring binding behavior of AR4A/AR3C IgG3C- and AR4A/AR3C CamX6, all complexes exhibited a 1:1 E1E2:Ab stoichiometry. This suggests that the longer hinges facilitate a favorable simultaneous interaction with both AR4A and AR3C epitopes on a single E1E2, indicating that these bsAbs utilize intra-E1E2 cross-linking as its main binding mechanism. The significantly lower abundance of unbound antibody molecules for the IgG3C-compared to the CamX6 format might suggest that the IgG3C-hinge allows for a more optimal angle of approach for intra-E1E2 cross-linking, resulting in the formation of more complexes, as we did not observe differences in binding strength for the two bsAbs in ELISA (Fig. 2E).

The peak distributions of observed E1E2:Ab complexes for the measured mAbs, bsAbs and cocktail are summarized in Figure 4E. This summary visualizes how hinge length of bsAbs can influence their binding modes, shifting from inter-E1E2 binding with shorter hinges to intra-E1E2 binding with longer hinges. Schematic representations of different observed binding modes and forms of avidity are shown in Figure 4F.

A schematic view based on an AlphaFold2 prediction of AR4A/AR3C IgG3C-, overlayed with the E1E2 crystal structure, illustrates the suggested intra-E1E2 binding mode of this bsAb (Fig. 4G). The flexibility of the open IgG3C-hinge allows the optimal angle of approach for cross-linking both epitopes on a single glycoprotein. In contrast, the more rigid hinge of an IgG1 bsAb restricts the Fabs from achieving the correct angles of approach for intra-E1E2 binding, instead favoring inter-E1E2 binding (Fig. 4H).

## Discussion

In this proof-of-concept study we present, to our knowledge, the first bsAbs designed to target different epitopes on the HCV E1E2 glycoprotein. We generated a panel of 60 bsAbs by combining 21 specificities from 8 HCV bNAbs. We show that HCV bsAbs can be successfully produced using CrossMab and cFAE technology, of which the former allows for hinge length modifications to provide various levels of Fab flexibility. All bsAb formats yielded properly folded antibodies, retaining their binding capacity, thermostability and affinity to FcγRIIIa.

We observed that the bsAbs exhibited a moderate increase in overall breadth compared to single mAbs, neutralizing the majority of tested viruses – sometimes more effectively than the parental mAbs and corresponding cocktails. Notably, the bsAbs could recover activity even when one arm was non-neutralizing as a single mAb. BsAbs may therefore be able to prevent viral escape when one of the arms is inactive against a specific strain or mutational variant, while also accommodating the wide variation in neutralization sensitivity across different tiers of HCV strains. In this scenario, one arm of the bsAb could be designed to specifically target a neutralization resistant HCV strain, while the other might cover a wider range of easier to neutralize viruses. This was particularly evident for one of the leading bsAb candidates identified in this study, AR4A/AR3C IgG3C-, which showed potency against a wide range of HCVpps and HCVccs with diverse sensitivities to antibody neutralization. In cases where AR3C was non- or very weakly neutralizing, such as with HCVpps AMS0230, AMS0232, and UKNP3.2.2, AR4A neutralization allowed this bsAb to restore its activity, and against UKNP3.2.2 neutralization was enhanced compared to the corresponding cocktail.

The neutralization screen of all bsAbs identified several promising candidates with improved potency, while also highlighting differences in the contribution of individual HCV mAbs to neutralization efficiency when incorporated into bsAb formats. For example, interesting differences were observed in bsAbs that paired AR4A with distinct bNAbs targeting the CD81bs. The angles of approach of bNAbs to the CD81bs can differ substantially (22, 52, 53) and this variation can be crucial when seeking synergistic epitope targeting and selecting the appropriate hinge to direct cooperative binding. For example, a bsAb combining AR4A with AR3C showed enhanced potency with the longer IgG3C-hinge, whereas the combination of AR4A with HEPC74, another CD81bs antibody but with a different angle of approach, performed better with the shorter IgG1 hinge or as a cocktail of two mAbs. The latter confirms results from a previous study, in which the AR4A and HEPC74 combination in a mAb cocktail was also identified as one that can provide synergy (29). Previous findings also showed that a mixture of AR4A-like and CD81bs-targeting antibodies is correlated with viral clearance in patients (16, 24), highlighting the clinical importance of examining the binding modes of these mAb classes in combination.

We also noticed a trend of improved activity for IgG1 hinge bsAbs containing AP33, with several demonstrating higher potencies than their corresponding cocktails. This improvement may result from a cooperative binding mechanism, where AP33 works in combination with other bNAbs, enhancing avidity by binding two epitopes on either the same E1E2 or neighboring E1E2s simultaneously. Another possibility is that one arm of the bsAb causes a conformational change in E1E2, improving the binding efficiency of the other arm. For example, AP33 binding could expose the CD81bs and thus enable AT1209 engagement, as has been proposed previously for similar mAbs HEPC98 and HEPC74 when used in combination (29). Previous studies have also reported enhanced binding of certain HCV bNAbs to E1E2 when another bNAb is already bound (43, 54), and the synergistic neutralization activity observed with mAb cocktails (28, 29) was shown to stem from distinct inhibition mechanisms of the individual components (29). Our results indicate that combining distinct specificities in bsAbs can provide additional cooperative benefits.

Bivalent binding is a crucial property of antibodies, as the simultaneous engagement of two epitopes is usually essential to achieve strong binding and effective neutralization (46, 55, 56). Typically, an IgG first attaches to an epitope with one Fab arm, and as the second Fab arm searches for another epitope, the first may detach and bind to an adjacent epitope. This dynamic process, known as “stochastic walking” on the surface of an antigen (57), makes accurately measuring and predicting avidity effects of bivalent (and bispecific) antibodies challenging. Several factors must be carefully considered, including the inherent flexibility of the hinge region, which influences the angle of approach for each Fab, and the vicinity of epitopes on an antigen or the density of antigens on a surface available for antibody cross-linking (58).

To gain deeper insights into the binding modes of our bsAbs, we used MP to determine the preferred binding stoichiometries of AR4A and AR3C, both when combined into bsAbs with different hinge lengths and as a cocktail. The IgG1 hinge bsAb and a cocktail formed higher-order complexes (1:2, 2:1, 2:2 and 3:2 E1E2:Ab) illustrating the diverse binding approaches, and potential avidity effects that can stem from combining different Fab specificities. However, the most striking observation was that both AR4A/AR3C bsAbs with long hinges strictly preferred 1:1 E1E2:Ab binding. Notably, AR4A/AR3C IgG3C-does so without any apparent loss of affinity or neutralization potency. In fact, this bsAb exhibited increased activity and a broad neutralizing response, suggesting that HCV mAbs may in some cases benefit more from intra-E1E2 avidity than from other binding modes. The calculated distance between AR4A and AR3C Fabs was found to be compatible with intra-E1E2 cross-linking when using a bsAb with a flexible open IgG3C-hinge. Conversely, the IgG1 hinge, although shorter, is more rigid, likely favoring inter-E1E2 cross-linking of the IgG1 hinge bsAb. Our results show that by strategically pairing different Fab arms and engineering the distance between them through hinge length modification, it is possible to achieve new angles of approach for E1E2 cross-linking. This can enhance avidity effects, allowing cooperative binding and consequently improve the neutralization activity of HCV bNAbs.

Testing the bsAbs in both HCVpp and HCVcc systems allowed us to obtain a more comprehensive view of their neutralizing breadth by capturing as much of the HCV genetic and antigenic diversity as possible. Previous studies have shown that structural differences between these two systems - such as the association of HCVcc, but not HCVpp, with apolipoproteins or differences in maturation of the glycoproteins, including variation in glycosylation profiles - can result in differences in neutralization sensitivity (59, 60). HCVpps are usually considered more sensitive to neutralization by serum antibodies and mAbs compared to HCVcc (44, 60). Although in this study different viral strains of HCVpp and HCVcc were used, previous studies have demonstrated sufficient correlation in neutralization sensitivity between the two, validating the use of both systems for assessing mAb neutralization activity (43, 60).

In our study, AR4A/AR3C IgG3C-ranked the highest against HCVcc, but not against HCVpp, where bsAbs with shorter hinges showed superior activity. Interestingly, a study using HIV bsAbs, which incorporated precisely designed linkers of different lengths between the two Fab arms and the Fc region, found that the optimal binding and neutralizing activity were achieved when bsAbs could target both epitopes on a single HIV envelope (Env) trimer at the same time (39). Given that the HIV virion is sparsely populated with trimers on its surface - a mechanism believed to help evade antibody neutralization - the study concluded that the ideal HIV-targeting (bs)Ab therapeutic would exhibit 1:1 Env:Ab binding. It is so far unknown how many functional and well-folded E1E2 complexes are present on an HCV virion (61). Our results suggest that HCVcc, which is closer to authentic HCV, may have fewer well-folded E1E2 copies on its surface compared to HCVpp, and that similarly to HIV, HCV might use a low envelope protein density as an antibody evasion mechanism. Alternatively, glycans or lipid molecules might shield E1E2 from neutralizing antibodies, giving bsAbs such as AR4A/AR3C IgG3C-, that utilize intra-E1E2 avidity as their primary binding mechanism, an advantage in inhibiting HCV. However, it is important to note that our MP analyses were conducted with soluble E1E2, thus our results may not fully reflect the dynamics of binding to membrane-bound E1E2 on the virion, where factors like steric hindrance might play a role.

It has recently been suggested that E1E2 could be expressed as a homodimeric complex on the viral surface (62), with “breathing” transitions from an open to a closed state, similarly to HIV Env. These transitions could further impact access to certain epitopes and enhance the benefits of certain avidity mechanisms. This underscores the importance of studying the binding mechanisms and potential synergistic effects of future mAbs and bsAbs against HCV E1E2.

Our results show that bsAbs can achieve cooperativity in simultaneous targeting of different HCV epitopes, resulting in enhanced neutralization potency and breadth. HCV bNAbs also hold the potential to be combined into tri- or tetra-specific antibodies or multispecific nanoparticles. By rationally designing antibody-like constructs with increased valency, it is possible to achieve enhanced avidity in binding and, most importantly, broaden virus neutralization (34, 35, 63, 64).

Future research into the structure of the HCV virion, the E1E2 glycoprotein heterodimer and the conformation(s) it adapts on the viral surface, as well as detailed studies of the binding mechanisms of HCV mAbs, will be crucial for designing new bivalent and multivalent antibodies. These constructs could potentially be used for (post-exposure) prophylaxis and/or treatment of HCV infection, in high-risk individuals or in cases of multi-DAA resistance. In parallel, bsAb constructs like the ones presented in this study, can serve as valuable tools for HCV research and immunogen design, aiding the development of a prophylactic vaccine.

This study provides new, generalizable avenues for improving HCV bNAbs and emphasizes the importance of achieving broad antiviral activity when designing next-generation antibody constructs against HCV.

## Acknowledgements

We thank Angela Schriek for providing the FcγRIIIa-dimer protein and helping with the setup of the ELISA. We thank Andreas Chrysostomou for help with the statistical ranking analysis. We thank Steven de Taeye and Mitch Brinkkemper for helpful discussions.

## Author contributions

Conceptualization: L.R., K.S. and J.S.

Methodology: L.R., A.O., T.K., I.Z., A.C., J.B., V.C.Y., J.M.G. and K.S.

Investigation: L.R., A.O., T.K., I.Z., J.C.P., S.K. and V.S.

Resources: L.R., A.O., J.C.P., F.M., A.C., J.B., A.J.R.H., J.M.G. and K.S.

Data curation and visualization: L.R. and T.K.

Supervision: A.O., J.B., M.J.vG., R.W.S., V.C.Y., A.J.R.H., J.M.G., K.S. and J.S.

Funding acquisition: J.B., M.J.vG., R.W.S., A.J.R.H., J.M.G., K.S. and J.S.

Writing-Original draft: L.R. and K.S.

Writing-Review and editing: all authors.

## Materials and methods

### Cell lines

FreeStyle human embryonic kidney (HEK) 293F cells (Life Technologies), a female human embryonic kidney cell line adapted to grow in suspension, were used for the production of recombinant proteins, including all antibody constructs in this study. Cells were cultured in FreeStyle 293 expression medium (Thermo Scientific) at 37°C with 8% CO_2_ and shaking at 125 rpm. For HCVpp production and neutralization assays, CD81KO HEK293T cells (ATCC, CRL-11268) and hepatocyte-derived carcinoma Huh-7 cells (gift from François-Loїc Cosset) were used, respectively. CD81KO HEK293 T cells were generated by following the Invitrogen TrueGuide sgRNA CRISPRMAX transfection protocol as previously described (65) and kept in Dulbecco’s modified Eagle medium (DMEM) (Sigma-Aldrich) supplemented with 10% fetal calf serum (FCS) and 1% penicillin-streptomycin (PS) at 37°C with 5% CO_2_. Huh7 cells were kept in DMEM supplemented with 10 % FCS, 0.1 % PS, 1 % nonessential amino acids and 1 % HEPES buffer at 37°C with 5% CO_2_. For HCVcc production and neutralization assays, the human hepatoma cell line Huh7.5 was used and cultured as described previously (66). Briefly, cells were kept in DMEM + 4500 mg/L glucose + GlutaMAX-I + Pyruvate (Gibco/ThermoFisher #31966-021) containing 10% heat-inactivated FCS (Sigma), penicillin 100 U/mL and streptomycin 100 µg/mL (Sigma/Merck), at 5% CO_2_ and 37°C.

### Constructs design

For the two bsAb formats with longer hinges (ie. IgG3C- and CamX6 hinge bsAbs) we used the CrossMab technology, which combines specific knob-in-hole mutations (S354C and T366W in the “knob” arm, and Y349C, T366S, L368A and Y407V in the “hole” arm) and a CH1-CL switch to ensure correct heavy chain dimerization and light chain pairing. This ensures the correct formation of the desired bsAb product following the co-transfection of different heavy and light chains (67, 68). For the generation of CrossMab bsAbs, variable heavy and light chain sequences of AR4A (38), AT1618, AT1209 (16), AP33 (69), AR3C (38), HEPC74 (22), HC84.26 (70) and IGH505 (71) were cloned into standard mammalian expression vectors containing the constant regions of the human IgG1, which included hinge and Fc sequence modifications as described in Suppl. Fig. 1 (Genscript Biotech). Additionally, LS mutations were introduced in the Fc region, known to increase half-life of antibody molecules (72, 73). For bsAb generation by controlled-Fab arm exchange, a quickchange site-directed mutagenesis kit (New England Biolabs) was used to introduce F405L and K409R mutations in the heavy chain plasmids of the IgGs. The codon-optimized soluble his-tagged HCV E2 was described before and is based on the genotype 1a H77 strain (Genbank: ABN11232.1 (amino acids 384–659) and GenBank: AF009606.1 (amino acids 660–715)) (74). For measuring stoichiometry of the bsAbs to sE1E2, we used an engineered StrepII-tagged version of the H77c-derived soluble native-like E1E2 as described by Guest and colleagues (75). The sequence of the FcγRIIIa-dimer (158V, high affinity polymorphic variant) was provided by Bruce Wines and cloned into a pcDNA3.1 expression plasmid (76).

### Expression and purification of antibodies and viral proteins

HCV monoclonal antibodies and bsAbs were produced as previously described (77). Briefly, for each antibody, the plasmids were co-transfected in a suspension of 125 mL of HEK293F cells, in a 1:1 HC:LC ratio for the monoclonal antibodies, including mAbs mutated for cFAE, and a 1:1.2:1:1.2 Fab HC/Fab LC/CrossFab HC/CrossFab LC ratio for the CrossMab bsAbs. After 5 days, the supernatants were filtered with 0.22 μm pore size SteriTop filters (Millipore), run over a column containing 0.5 mL of protein G beads (Pierce), and washed three times with PBS. The antibodies were eluted with 18 mL elution buffer (0.1 M glycine pH 2.5) into 2 ml of neutralization buffer (1 M TRIS pH 8.7). They were then concentrated and buffer exchanged to PBS using a 100 kDa VivaSpin20 column (brand). Antibody concentrations were measured using a NanoDrop 2000 (ThermoFisher). E2 and E1E2 proteins were purified by NiNTA or StrepTactinXT purification, respectively, as previously described (78). The FcγRIIIa-dimer protein was produced as previously described (76).

### Generation of bsAbs by controlled Fab-arm exchange (cFAE)

BsAbs containing a “normal” IgG1 hinge were produced using a controlled Fab-arm exchange protocol, as previously described (32). In short, equimolar amounts of antibodies containing either the F405L or K409K mutation were mixed with a solution of 2-mercaptoethylamine (2-MEA, 750 mM, pH 7.4; Sigma) and PBS and incubated for 5h at 31°C. Following the incubation, the 2-MEA was removed by washing and buffer exchange to PBS using 100kDa Vivaspin6 columns (Sartorius), and left overnight to enable successful reoxidation of the disulfide bonds between antibody chains.

### SDS-PAGE

For SDS-PAGE analyses 4 µL of antibody (1 mg/mL) was mixed with loading dye (25 mM Tris, 192 mM Glycine, 20% v/v glycerol, 4% m/v SDS, 0.1% v/v bromophenol blue in mQ water), incubated at 95 °C on a thermoblock for 10 min and loaded on a 10–20% Tris-Glycine gel (Invitrogen). For reducing SDS-PAGE, 1.2 µL of dithiothreitol (DTT; 100 mM) was added to each sample before heating. Gels were run in running buffer (25 mM Tris, 192 mM glycine, 0.5% SDS in distilled water) for 45 min at 200V at 4 °C. Gel staining was performed using PageBlue Protein Staining Solution (Thermo Fisher Scientific) overnight, followed by destaining with mQ water.

### Differential Scanning Fluorimetry (DSF)

Protein thermostability was evaluated with a Rotor-Gene Q instrument (QIAGEN). A fresh 1:200 mix of Sypro Orange dye (Sigma) was prepared in PBS, and 15 µL of dye was mixed with 10 µL of antibody (1 mg/mL). The intrinsic fluorescence signal was measured while temperature was increased from 40 to 95°C by 1°C every 5 seconds. The melting temperatures (Tm) were determined using the Rotor-Gene Q series software.

### Enzyme-linked immunosorbent assay (ELISA)

To determine bispecificity of bsAbs his-tagged HCV E2 (3 μg/mL in Tris-buffered saline (TBS)) was immobilized on NiNTA 96-well plates (Qiagen) for 2h at RT, followed by the monospecific, bispecific or a 1:1 cocktail of parental monospecific antibodies (10 μg/mL in casein) for 1.5h at RT. Next, anti-kappa LC or anti-lambda LC secondary antibodies (Bethyl Laboratories) were added (1:10000 in casein) and left to incubate for 45 min at RT. Plates were washed three times with TBS in between all incubation steps. After the last step, the plates were washed five times with TBS/0.05% Tween-20, followed by signal development and readout.

To test FcγRIIIa binding, we performed StreptactinXT ELISA. Soluble E1E2 protein (1 μg/mL in TBS) was coated on 96-well Strep-TactinXT coated microplates (IBA LifeSciences) for 2h at 37°C, followed by four washes in TBS + 0.05% Tween-20. Wells were blocked with assay buffer (1% BSA, 0.05% Tween-20, 1mM EDTA in PBS) for 1h at 37°C. Next, antibodies were 2-fold serially diluted in casein (Thermo Fisher Scientific), starting at 5 μg/mL and incubated for 1h at 37°C, followed by biotinylated FcγRIIIa-dimer protein (0.5 μg/mL in assay buffer) for 1h at 37°C and high sensitivity streptavidin-HRP detection antibody (1:2000 in assay buffer) for 1h at 37°C. In between every incubation step, plates were washed four times with TBS + 0.05% Tween-20. The same assay was performed without the addition of FcγRIIIa-dimer in parallel to assess potential differences in antibody:E1E2 binding. All experimental parameters were kept the same, except that primary antibodies were serially diluted 5-fold and followed by incubation with a 1:3000 dilution of HRP-labeled goat anti-human IgG (Jackson Immunoresearch) in casein for 45 min at 37°C. For all ELISA experiments signal development was performed by coating with a solution of 1% 3,3’,5,5’-tetramethylbenzidine (Sigma-Aldrich), 0.01% hydrogen peroxide, 100 mM sodium acetate and 100 mM citric acid for 5 min, followed by 0.8 M sulfuric acid to end the development reaction. Readouts of optical densities (OD) at 450 nm were performed using a spectrophotometer (BMG Labtech).

### Biolayer interferometry (BLI)

Antibodies were loaded onto a protein A biosensor (Sartorius) for 300 s with a concentration of 10 μg/mL, followed by a baseline in running buffer (PBS, 0.02% Tween, 0.1% bovine serum albumin (BSA)) for 60 s. The sensors were then immersed in a solution of E1E2 protein (150 mM) to measure association for 300 s, followed by a 300 s dissociation step in a new well containing running buffer. Antibodies and E1E2 protein were diluted in running buffer, and a well containing only running buffer was used to measure background binding. Sensors were regenerated between measurements using a buffer containing 10 mM glycine in PBS. BLI experiments were performed using an Octet K2 instrument (ForteBio), at 30°C and with agitation speed of 1000 rpm.

### HCV pseudoparticle production

HCVpp production was performed as described previously (77). Briefly, 1.5 × 10^6^ CD81KO HEK293T cells were seeded on a 10 cm^2^ dish one day before transfection. Cells were co-transfected with three plasmids: E1E2, an MLV Gag-Pol packaging construct and firefly luciferase (79). E1E2 constructs were based on the HCV glycoprotein sequences from the following strains (GenBank ID; mutation(s)): H77 (AAB67037; R564C, V566A, G650E), AMS0229 (OL855834.1), AMS0230 (OL855838.1), AMS0232 (OL855837.1; I442F), Ams1b (KR094962.1), AMS2b (KR094963.1), AMS3a (KR094964.1), Ams4d (ON623878), UKNP2.2.1 (KU285211.1), UKNP2.4.1 (KU285213.1), UKNP3.2.2 (KU285219.1), UKNP4.1.1 (ALV85530.1), UKNP5.2.1 (ALV85536.1) and JFH1 (KF700370.1). Plasmids were used in previously optimized ratios with a total amount of 6 µg of DNA and 12 µl of Lipofectamine 2000 (Invitrogen) in Opti-MEM (ThermoFisher). The following day, Opti-MEM was replaced by DMEM (Gibco)/10% FCS/0.1% PS. Supernatants containing the HCVpps were collected 48 h after transfection, sterile filtered through a 0.45 μm syringe filter and frozen at −80°C for long-term storage or 4°C when used within a week.

### HCVpp neutralization assay

HCVpp neutralization assays were performed as described previously. In short, Huh-7 cells were seeded at a density of 15 × 10^3^ per well in 96-well culture plates in Huh-7 culture medium (DMEM/10% FCS/0.1% PS/ 1% nonessential amino acids/1% HEPES buffer). After 24h, HCVpps were incubated with antibodies in triplicate, either with a specific concentration of antibody or with 3 or 4-fold serially diluted antibody, at 37 °C in 5% CO_2_. After a 1 h incubation at 37°C in 5% CO_2_, the HCVpp/antibody mixtures were added to the Huh-7 cells. Following a 4 h incubation at 37°C in 5% CO_2_, Huh-7 media was added to the cells and the plates were left to incubate for 72h. Afterwards, cells were lysed and luciferase activity was measured using the Nano-Glo Luciferase Assay System (Promega) with a Glomax plate reader (Turner BioSystems).

### Production of HCVcc virus stocks

HCVcc virus stocks were generated as described previously (44). Briefly, Huh7.5 cells cultured in DMEM + 4500 mg/L glucose + GlutaMAX-I + Pyruvate/10%FCS/ penicillin 100 U/mL and streptomycin 100 µg/mL were infected the day after seeding with JFH1-based recombinant viruses harboring Core-NS2 of the following genotypes (strains) with the indicated cell-culture adaptive mutations (amino acid numbers according to the H77 reference sequence, genbank accession number AF009606): 1a(TN) with R1408W (80), 2a(J6) (81), 3a(S52) with I793S and K1404Q (66), 4a(ED43) with T827A and T977S (82), and 5a(SA13) with A1022G and K1119R (83). Cultures were split and supernatants were harvested every 2–3 days. Supernatant from timepoints with the highest infectivity titers determined as described previously (84), were pooled and HCVcc virus stocks were stored at −80°C. If needed, virus stocks were concentrated using Amicon 100 kDa centrifugation filters (Millipore). HCVcc virus stock E1E2-sequences were determined by Sanger sequencing (Macrogen) (85) and were identical to the plasmid sequences reported for each recombinant.

### HCV cc neutralization assay

In vitro cell-based HCVcc neutralization assays were carried out as described previously with slight modifications (83). In brief, Huh7.5 cells were plated at 9000 cells/well in clear-bottom 96-well plates (Nunc) the day before the experiment and incubated overnight at 37 °C and 5% CO_2_. MAbs, bsAbs or cocktails of mAbs mixed 1:1 were diluted in culture medium (DMEM t+ 4500 mg/L glucose + GlutaMAX-I + Pyruvate/10%FCS/penicillin 100 U/mL and streptomycin 100 µg/mL) and mixed with HCVcc, diluted to yield focus forming unit (FFU) counts of 30–200 in the absence of antibodies as determined in pilot experiments, and left to incubate 1 h at 37 °C and 5% CO_2_. Experimental wells were set up in triplicates. The C211 antibody (86) and SA13/JFH1core-NS5B virus (84) were used as neutralization controls. Six blank wells with neither antibody nor virus, and six wells with untreated virus were included in each plate. The antibody/virus mixes were added to the plated cells, 100 µl/well, and incubated at 37 °C and 5% CO_2_ for 3h. The cells were then washed with pre-warmed PBS and fresh culture media was added. The plates were left to incubate for 48+/-2h after infection at 37 °C and 5% CO_2_, and subsequently fixed with ice-cold methanol. The plates were incubated with 3% H_2_O_2_ for 10 min at RT and immunostained with primary anti-NS5A antibody 9E10 (81) at 1:5000 dilution in PBS (Sigma) with 1% BSA (Roche) and 0.2% skimmed-milk (Easis) at 4 °C overnight, followed by incubation with secondary antibody ECL anti-mouse IgG horseradish peroxidase (HRP)-linked whole antibody (GE Healthcare Amersham) at 1:500 in PBS +0.1% Tween-20 for 2h at RT. The Pierce DAB Substrate Kit (Thermo Scientific) was used according to the manufacturer’s instructions for visualization of FFU. FFU were counted automatically using an ImmunoSpot series 5 UV analyzer (CTL Europe GmbH) with customized software as described previously (87). The % neutralization were calculated by relating FFU counts of experimental wells to the mean FFU count of untreated virus wells.

### Mass photometry

Mass Photometry (MP) experiments were performed on a OneMP mass photometer (Refeyn Ltd., UK). The analysis was performed on sample carrier slides prepared in-house from microscope coverslips (24 mm × 50 mm; Paul Marienfeld GmbH) and reusable cell culture gaskets (Grace Biolabs). The data were recorded using vendor provided software AcquireMP (Refeyn Ltd., UK), for 60 s with a scan rate of 100 frames per second using the Medium field of view. Antigen-antibody complexes were analyzed as previously described (49, 88). Briefly, soluble E1E2 and antibody were mixed in a 3:1 molar ratio, resulting in 500 nM and 166.7 nM E1E2 and antibody, respectively. The samples were incubated (30 min, 30°C) and analyzed in 10 nM final concentration (for E1E2). All experiments were conducted in PBS, pH = 7.4. The mass calibration was performed using an in-house prepared standard protein mixture (IgG-halfbody 73 kDa; IgG 149 kDa; apoferritin 513 kDa; IgM 1006 kDa). Calibrated events were exported and processed by an in-house developed Python pipeline (88) using NumPy (89), pandas (90), Matplotlib (91), SciPy (92) and seaborn (93). Mass histograms were plotted with 4 kDa bin widths and resulting mass distributions were annotated manually.

### Structure visualization and prediction

Protein structures, complexes and predictions were visualized and distance between atoms was calculated with UCSF ChimeraX 1.8, developed by the Resource for Biocomputing, Visualization, and Informatics at the University of California, San Francisco, with support from National Institutes of Health R01-GM129325 and the Office of Cyber Infrastructure and Computational Biology, National Institute of Allergy and Infectious Diseases (94). For the prediction of AR4A/AR3C IgG3C-structure, AlphaFold-2.3.1 multimer was used (95, 96). Code available at: https://github.com/google-deepmind/alphafold.

### Quantification and statistical analysis

All midpoint mAb inhibition concentrations (IC_50_ values) were determined, and data visualization and statistical analyses were performed in GraphPad Prism 8.3.0. Where applicable, statistical parameters are reported in the figure legends. Statistical analyses of bsAb ranking reported in Supplemental Fig. 4B were performed in Microsoft Excel as follows. The seven bsAb candidates (listed in Fig. 3C) were assigned rank scores (1–7) for each of the the following 4 properties:

a= rank of bsAb by IC_50_per tested virus
b= rank of bsAb by IC_50_ ratio bsAb/cocktail per tested virus
c= rank of bsAb by IC_50_ ratio bsAb/more potent parental mAb per tested virus d= rank of bsAb by IC_50_ ratio bsAb/less potent parental mAb per tested virus.

A total ranking score (A) per bsAb for all viruses combined (calculated separately for the HCVpps and HCVccs tested) was calculated as follows:

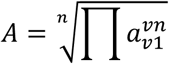

where v=virus (HCVpp or HCVcc), n=number of tested viruses (4 for HCVpp, 5 for HCVcc). This was repeated for each of the 4 properties, resulting in total ranking scores A, B, C and D for each bsAb. Total ranking scores (A-D) were then assigned a weight factor according to importance and relevance to our study, for a final ranking value per bsAb calculated as follows:

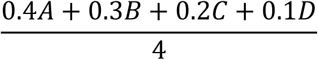

Final ranking values were sorted from lowest to highest for a total final ranking score of the bsAbs.

---

**Supplemental figure 1.**
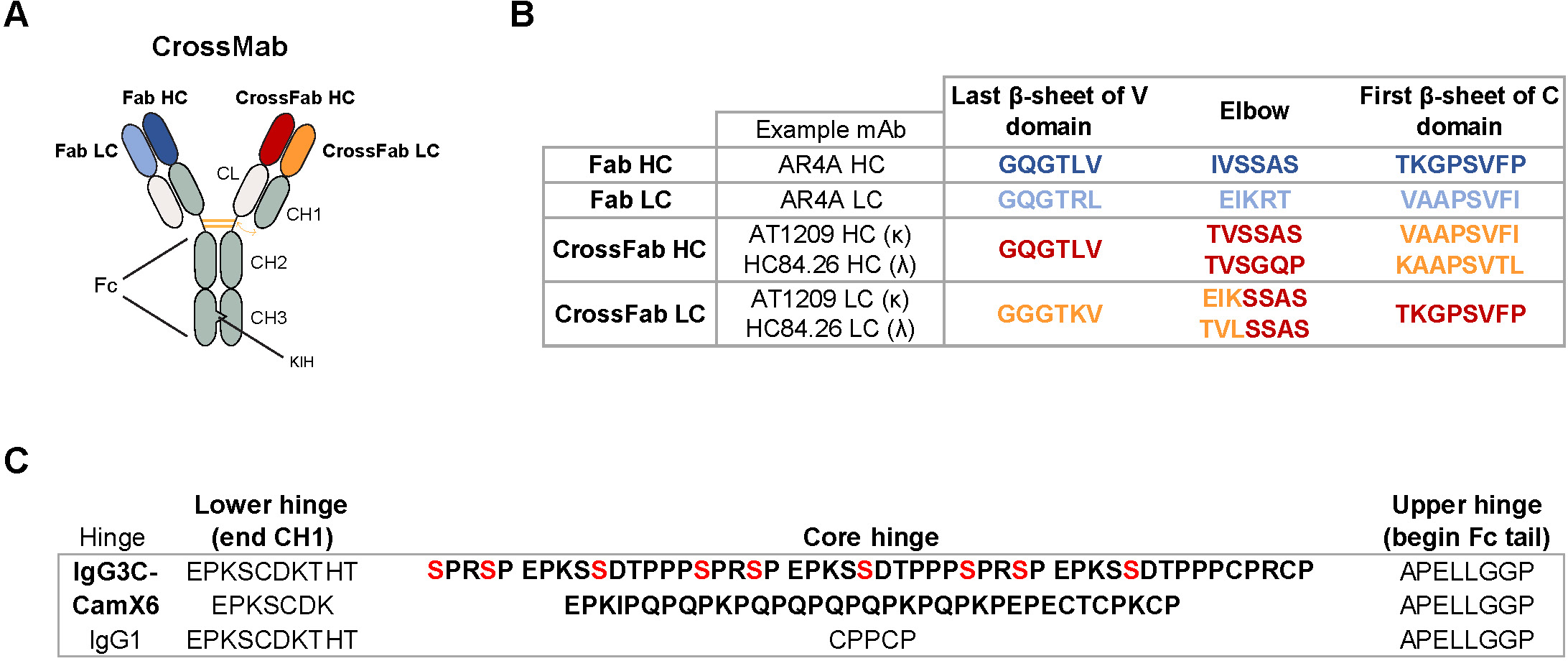
CrossMab bsAb design. **A** Schematic representation of a CrossMab bsAb showing structural components involved in bispecificity. **B** Sequences of elbow and adjacent regions of HC and LC in Fab and CrossFab constructs for CrossMab production. Two examples of CrossFab designs are provided, one for the kappa LC isotype (AT1209) and another for the lambda LC isotype (HC84.26). The same colors are used for the different Fabs as in A. **C** Sequences of the hinge regions in the bsAb constructs used in this study, in comparison to the natural IgG1 hinge. The IgG3C-hinge originates from the natural IgG3 hinge, with all cysteine residues replaced by serines, except for the last two (positions indicated in red). HC, heavy chain; LC, light chain; CH1, constant heavy chain region 1; CL, constant light chain region; KIH, knob-in-hole.

**Supplemental figure 2.**
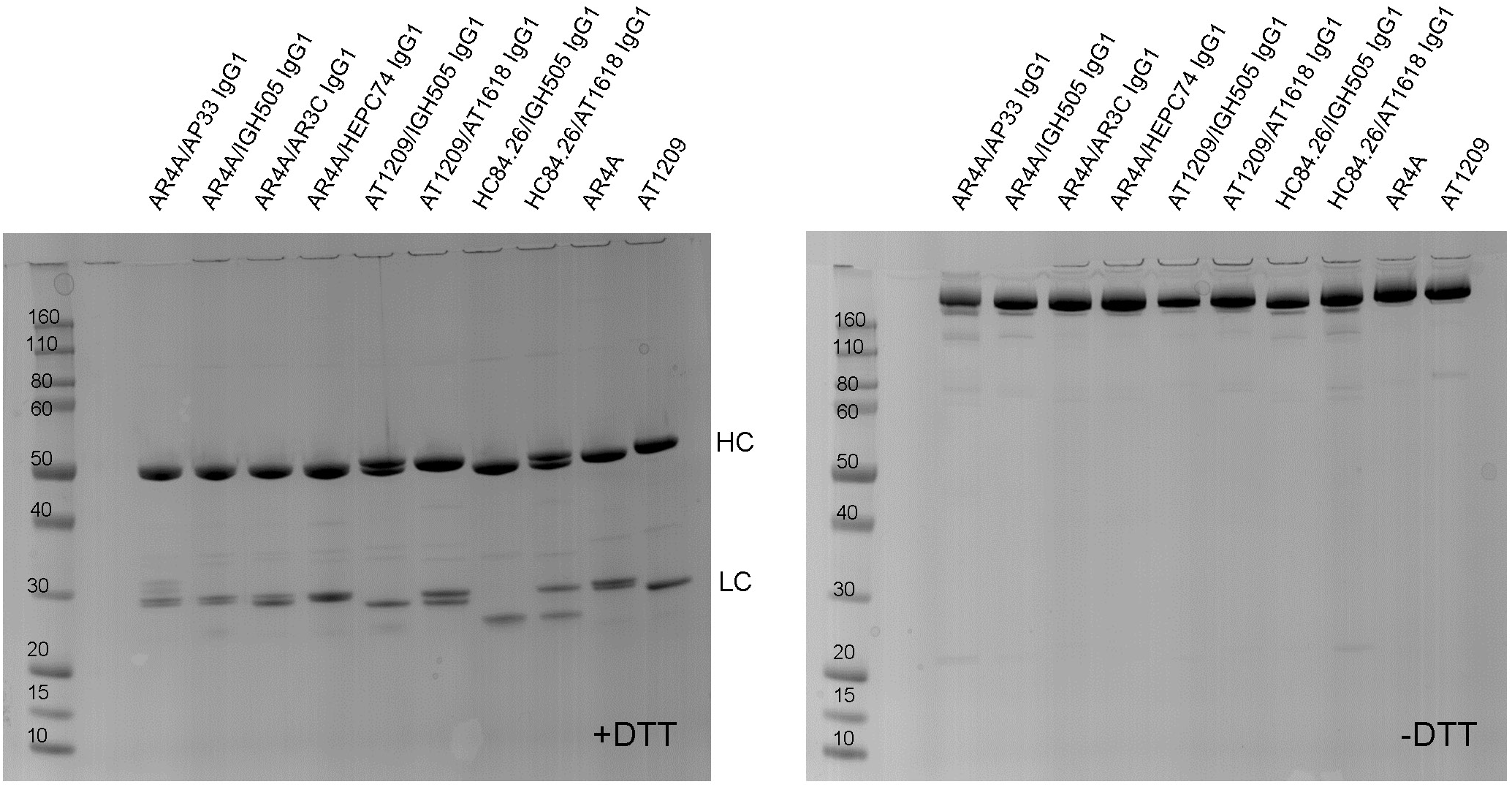
IgG1 hinge bsAbs produced by cFAE. SDS-PAGE analysis under reducing (left) and non-reducing (right) conditions of representative bsAbs produced by controlled Fab-arm exchange, containing a standard IgG1 hinge region. AR4A and AT1209 mAbs were included as controls.

**Supplemental figure 3.**
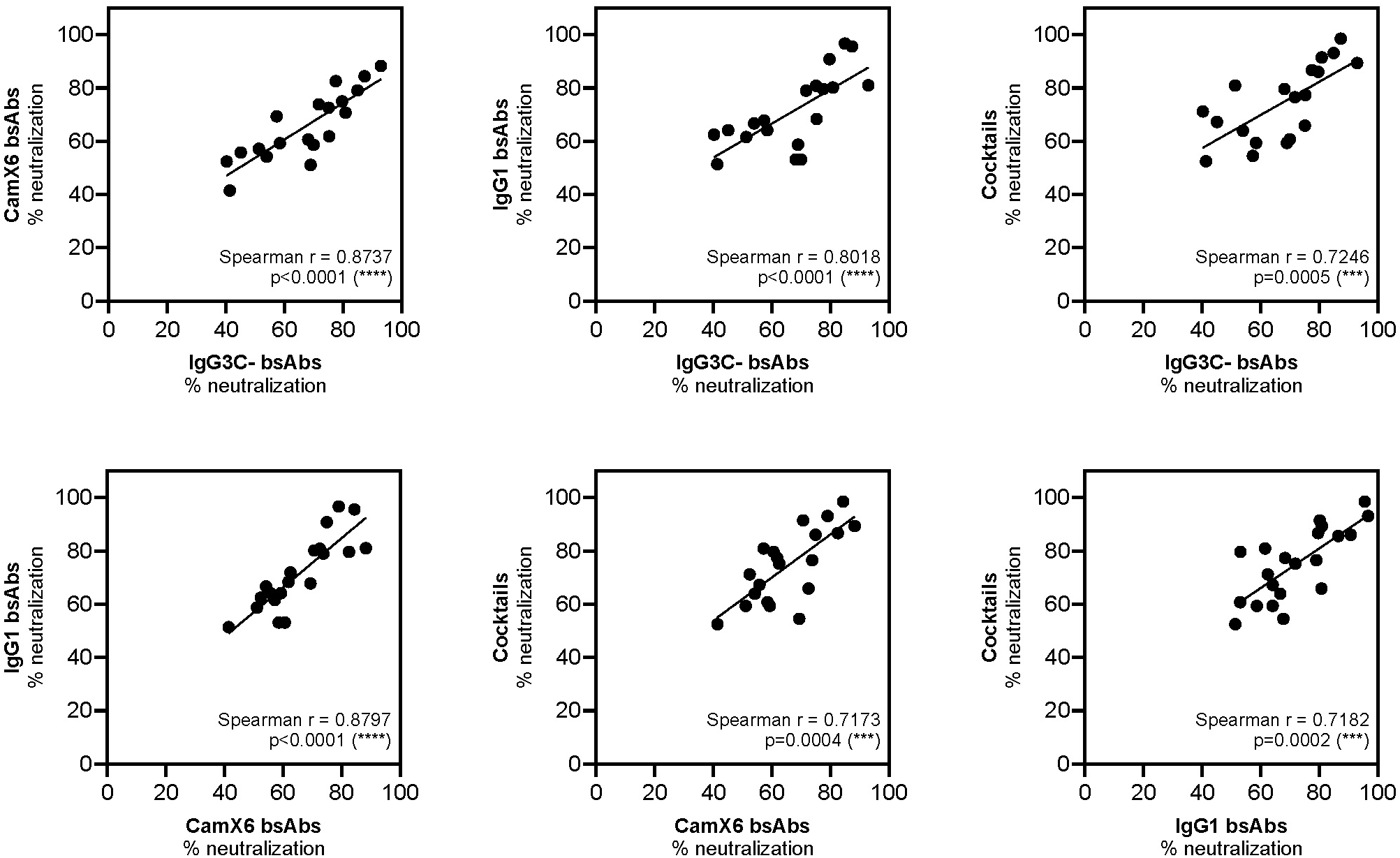
Correlation of HCVpp neutralization by different bsAb formats and antibody cocktails. Related to Fig. 3B. Shown are the correlations between the percentage of neutralization achieved by the different groups of antibodies tested. Spearman correlation (r) and p values were calculated using GraphPad prism 8.3.0.

**Supplemental figure 4.**
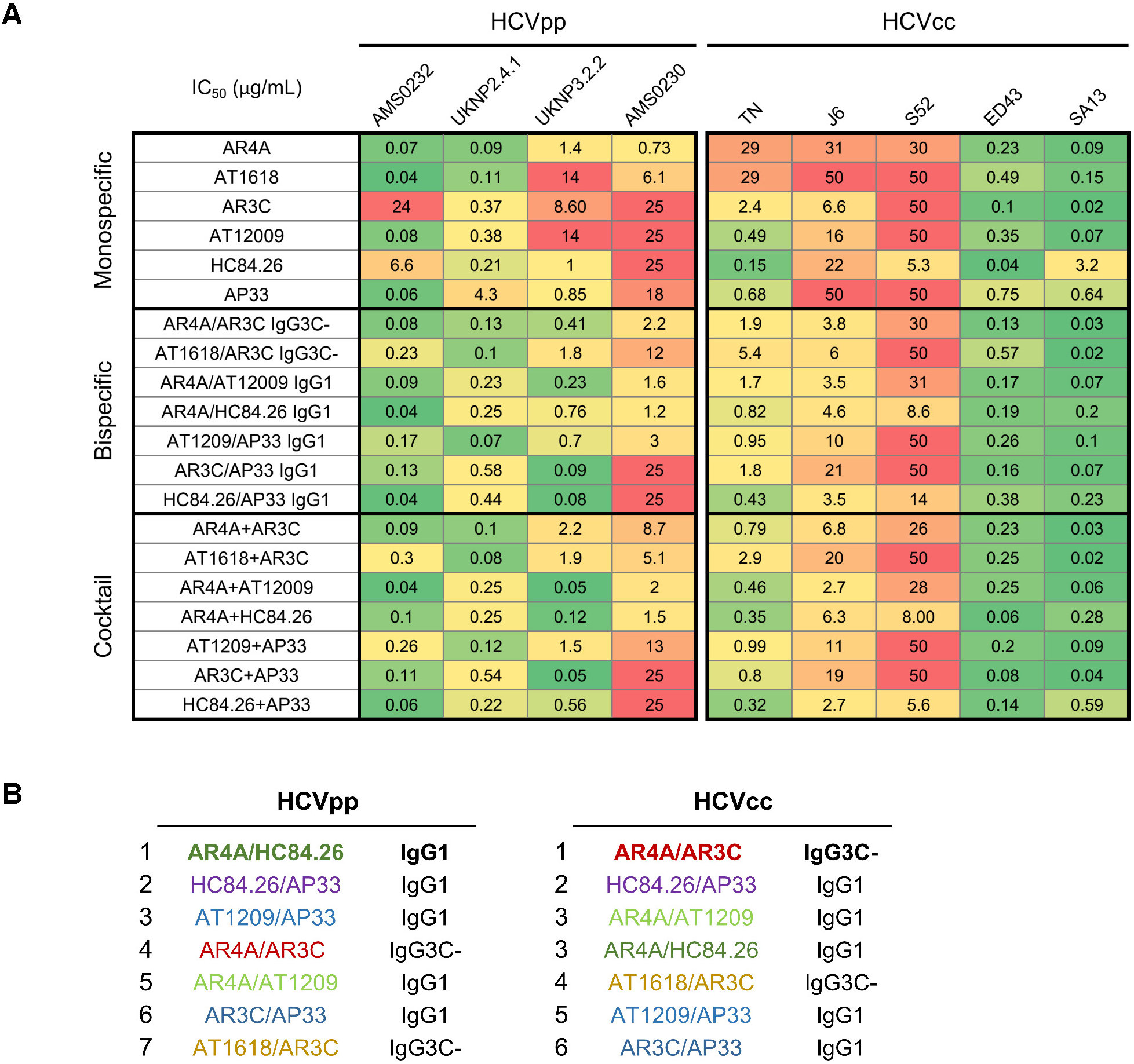
HCVpp and HCVcc neutralization by bsAbs and control antibodies. Related to Fig. 3D, E and F. **A** Half-maximal inhibitory (IC_50_) values (µg/mL) for neutralization of HCVpp and HCVcc by E1E2-targeting mAbs, selected bsAbs and corresponding cocktails. Each IC_50_ value represents the mean of at least two independent experiments performed in triplicate for HCVpp or one experiment performed in triplicate for HCVcc. IC_50_ values exceeding 25 µg/mL (HCVpps) or 50 µg/mL (HCVcc) were rounded down to 25 or 50, respectively, and classified as non-neutralizing. **B** Final ranking of seven selected antibodies (as listed in Fig. 3C) based on IC_50_ values and the ratio comparisons between bsAbs and cocktail, as well as bsAbs and their more or less potent parental antibodies. The analysis was performed separately for the HCVpp and HCVcc strains. Notably, AR4A/AT1209 and AR4A/HC84.26 IgG1 hinge bsAbs share identical rankings against HCVcc. Details of the analysis are described in the methods section under Quantification and Statistical Analyses.

**Supplemental figure 5.**
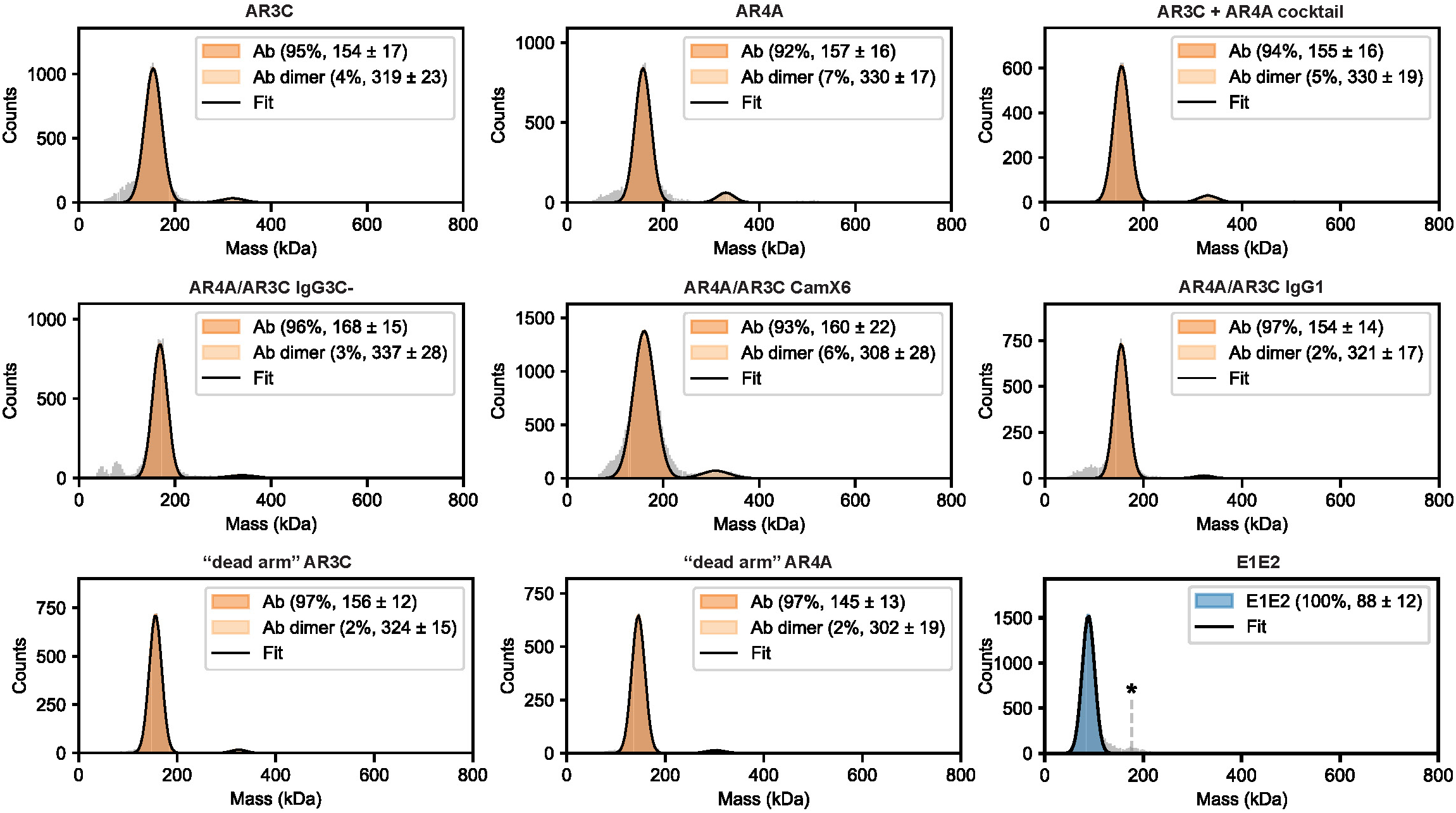
Mass photometry histograms of antibodies and E1E2 protein. A predominantly single mass distribution was observed for all antibodies, with expected masses of approximately 150kDa for IgG1 constructs, 160 kDa for the bsAb containing a CamX6 hinge, and 168 kDa for the bsAb with an IgG3C-hinge. Antibody dimers were detected at levels below 7% for both monospecific and bispecific antibodies, indicating minimal aggregation during Ab purification, unrelated to bispecific antibody production. For the soluble E1E2 protein, a major distribution was observed around 88 kDa, with a minor peak in the 150-200 kDa range, likely corresponding to an E1E2 dimer, annotated by an asterisk.

